# Preparing high-quality chromosome spreads from *Crocus* species for karyotyping and FISH

**DOI:** 10.1101/2024.11.18.624136

**Authors:** Abdullah El-nagish, Susan Liedtke, Sarah Breitenbach, Tony Heitkam

**Affiliations:** Faculty of Biology, Institute of Botany, Technische Universität Dresden, 01069 Dresden, Germany; Botany and Microbiology Department, Faculty of Science, Sohag University, 82524 Sohag, Egypt; Department of Biology, Institute of Biology I, RWTH Aachen University, Worringer Weg 3, 52074 Aachen, Germany

**Keywords:** Hydroxyurea-Colchicine Method, Nitrous Oxide, Hydroxyquinoline, Ice Water pretreatment, Metaphase Index, saffron, *Crocus sativus*, fluorescent *in situ* hybridization

## Abstract

**Background:** The saffron-producing *Crocus sativus* and its wild relative *C. cartwrightianus* are key species for understanding genetic evolution in this genus. Molecular-cytogenetic methods, especially fluorescent *in situ* hybridization (FISH), are essential for exploring the genetic relationships in this genus. Yet, preparing high-quality chromosomes for FISH analysis across *Crocus* species remains difficult. A standardized protocol for achieving clear and well-separated mitotic chromosomes is still lacking. This study assesses the effectiveness of four chromosome fixation methods for optimal chromosome spread preparation in *Crocus*. Root tips of different *Crocus* species were treated with four chromosome preparation methods namely hydroxyurea-colchicine (HC), nitrous oxide (NO), hydroxyquinoline (HQ), and ice water (IW) pretreatments to investigate their effectiveness in producing high-quality mitotic chromosome spreads. Metaphases obtained by the four methods were analyzed to assess their quality and metaphase index.

**Results:** Evaluation of 22,507 nuclei allowed us to confidently recommend a protocol for *Crocus* chromosome preparation. Among the methods, ice water pretreatment yielded the highest metaphase index (2.05%), more than doubling the results of HC (1.08%), NO (1.15%), and HQ (1.16%). Ice water-treated chromosomes exhibited better chromosome morphology, with relatively proper size, and non-overlapping chromosomes that were optimal for FISH analysis. Ice water pretreatment was also applied to *C. cartwrightianus*, the diploid progenitor of *C. sativus*, where it demonstrated similar efficacy. DAPI staining of chromosomes in both species allowed for clear visualization of intercalary and terminal heterochromatin. FISH analysis using 18S-5.8S-25S and 5S rDNA probes confirmed the utility of IW-prepared chromosome spreads for cytogenetic studies.

**Conclusions:** We strongly recommend ice water pretreatment as a suitable and effective method for obtaining many metaphase spreads of high-quality in *C. sativus* and related species, particularly for applications involving a detailed cytogenetic analysis.

## Background

The genus *Crocus* (Iridaceae) comprises around 250 species widely distributed over a wide range of climatic areas [1,2] and is known for its variable chromosome numbers [3-5]. Among the species of *Crocus*, only *C. sativus* is used as a crop and thus receives the most attention. *C. sativus* is the source of saffron, one of the highest priced spices of the world, processed from dried stigmas of manually harvested flowers. It is a cash crop for agriculture-based communities living off marginal areas in Iran, North Africa, countries surrounding the Mediterranean basin, and Kashmir. Despite its economic relevance, we are just beginning to understand the genomic and chromosomal constitution of saffron crocus and related species.

*C. sativus* is a male-sterile triploid species harboring eight chromosome triplets (2n = 3x = 24) and having a genome size of 1C = 3.45 Gbp [6]. Due to its triploidy, saffron crocuses can only be propagated vegetatively. As all saffron accessions around the globe have a similar genome, it is generally accepted that triploid saffron emerged only once [3, 7-11]. We and other groups recently showed that cytotypes of *Crocus cartwrightianus* have been the sole precursors of saffron’s triploidy, and that its emergence can be traced to the Aegean Bronze Age in Greece [3,4,12,13]. *C. cartwrightianus* is a diploid species (2n = 16) with high genetic diversity [3,4,12]. Nevertheless, despite these recent insights into the origin of saffron crocus, the *C. cartwrightianus* cytotypes that may enable targeted re-breeding and improvement of saffron traits have not yet been identified. Similarly, the origin of the individual chromosomes within saffron’s chromosome triplets is still unclear, especially as one chromosome is heteromorphic [14]. However, genomic and cytogenetic analyses in the genus *Crocus* may provide detailed insights into the chromosome structure of *Crocus* species, but robust and widely applicable protocols are still lacking.

Fluorescent *in situ* hybridization (FISH) is a powerful cytogenetic technique to study structure and function of chromosomes, polyploidy and genome evolution. In particular, the physical mapping of tandemly repeated DNA sequences provides informative cytogenetic landmarks for unequivocal chromosome identification in many plant species [15-19]. The first *in situ* hybridizations along *Crocus* chromosomes already showed the potential of repeat probes in this genus, yielding a range of distinct signals and allowing first chromosome assignments [20-22]. Recently, we developed a karyotyping mix for *Crocus* species that is composed of six tandem repeat probes [4] and that opens the *Crocus* genus for comprehensive molecular-cytogenetic analyses to clarify the genetic details of saffron crocus’ ancestry. However, streamlining FISH analysis across a range of species, cytoypes and accessions requires a robust protocol for properly dispersed mitotic chromosomes for its application. Despite being studied cytogenetically for several decades, species of the *Crocus* genus still remain challenging targets for chromosome preparation. Usually, due to the strict annual growth and the small size of the plant, material is limited, especially of wild accessions. In addition, as the chromosomes are large, some preparation methods such as dropping [23-26] are not recommended. Therefore, a comparative study testing different methods to obtain high-quality mitotic chromosomes from *Crocus* species is needed.

Here, using the crop plant *C. sativus* for its high economic value and *C. cartwrightianus* for its scientific interest, we compared the effectiveness of four chromosome fixation methods for obtaining *Crocus* chromosome spreads. We use (1) the hydroxyurea-colchicine method, (2) the nitrous oxide method, (3) the hydroxyquinoline method, and (4) ice water treatments and analyze them for their yield in obtaining properly spread mitotic chromosomes and mitotic index.

## Methods

### Plant materials and time of harvest

We used root tips of *C. sativus* (corms collected commercially), *C. cartwrightianus HKEP 1517* and *C. cartwrightianus* (Attica S FB19-63 (2)) was provided by F. Blattner and D. Harpke (IPK Gatersleben, Gatersleben, Germany). All plants were grown under glasshouse conditions. Root tips were collected in the early morning hours (07.00-08.00).

### Experimental design

Root tips of *C. sativus* were treated with four different synchronization methods. Each method was repeated six times:

1. <u>H</u>ydroxyurea-<u>c</u>olchicine method (HC): Corms with roots were incubated for 18 h in liquid, 0.5x LM medium containing 1.25 mM hydroxyurea. The material was kept in 125 mL Erlenmeyer flasks placed on an orbital shaker at 150 rpm at room temperature. After three rinses with 0.5x LM medium without hydroxyurea, the material was incubated for 6 h in fresh medium followed by treatment with the medium containing 0.6 % (w/v) colchicine for 20 h [27]. Treated root tips were excised and fixed in a 3:1 (v/v) ethanol: acetic acid solution for 24h at 4 °C.
2. <u>N</u>itrous <u>O</u>xide method (NO): Root tips were incubated in a pressure-tolerant cylinder, with nitrous oxide gas applied for 45 min at room temperature at 10 bar [28]. After this, the root tips were fixed in a 3:1 (v/v) methanol: acetic acid solution for 24h at 4 °C [29].
3. <u>H</u>ydroxy<u>q</u>uinoline method (HQ): Roots from individual corms were collected, synchronized in 2 mM hydroxyquinoline for 5h and fixed in a 3:1 methanol: acetic acid [4].
4. Ice <u>w</u>ater method (IW): Corms with roots were placed in container filled with ice water, kept inside a refrigerator at 4° C for 18 hours. Roots of 2-3 cm long were cut from and were fixed in methanol: acetic acid (3: 1) for 2h at 4 °C, fresh fixative was add and kept for 24h at 4 °C.

### Protocol of chromosome preparation from *Crocus* material

#### Enzyme treatment of root tips

1. Wash root tips 1x in dist. H_2_O for 5-10 min, 2x in 4 mM citrate buffer, pH 4.5 for 5 min each.

2. Dissect meristematic tips using a sharp scalpel and transfer them into a petri dish with 20–30 μl of enzyme mixture. The enzyme solution consists of 2% (w/v) cellulose from *Aspergillus niger*, 4% (w/v) cellulase ‘Onozuka R10’ from *A. niger*, 2% (w/v) hemicellulase from *A. niger*, 0.5% (w/v) pectolyase from *Aspergillus japonicus* and 20% (v/v) pectinase from *A. niger* in citrate buffer (4 mm citric acid and 6 mm sodium citrate).

3. Incubate at 37°C for 2.0-2.5 h depending on species (Table 1).

**Table 1:** Incubation times of different *Crocus* root tips for all chromosome preparation protocols.

| Species /Accession | Incubation time<br>in enzyme<br>mixture | Incubation time<br>on hot plate |
| --- | --- | --- |
| *C. sativus* | 2.30 h | 60 sec |
| *C. cartwrightianus* HKEP 1517 | 2.00 h | 30 sec |
| *C. cartwrightianus* (Attica S FB19-63 (2)) | 2.15 h | 30 sec |

#### Spread preparation

1. Single root tips were transferred onto slides, macerated in 50-60 μl of 45% acetic acid with needles for 120 sec. 2. Add an extra drop of 45% acetic acid, mix with a needle, spread on a hot surface at 55oC for 30-60 sec depending on species (Table 1).

#### Spread fixation

1. Surround with drops of freshly prepared fixative (3:1 methanol:acetic acid), dropwise fresh fixative on top of the slide as well.
2. Let the fixative run down, rinse with more fixative.
3. Air-dry the slides.
4. Store slides in a Coplin jar in a freezer until usage.

### Assessing the quality of the mitotic chromosome spreads via phase contrast and fluorescent microscopy

The chromosome preparations were assessed via phase contrast and fluorescent microscopy. For the latter, mitotic chromosome spreads were prepared according to the protocol above. Slides were equilibrated in 4x SSC / 0.2 % Tween 20 for 5 min at 37 °C. Excess liquid was carefully removed, then, 20 μl DAPI-Citifluor AF1 were added on each slide and covered with glass cover.

Wide-field imaging was performed with a Zeiss Axioimager M1 UV epifluorescence microscope with appropriate filters, and equipped with an ASI BV300-20A camera coupled with the Applied Spectral Imaging software (Applied Spectral Imaging, Carlsbad, CA, USA). The images were processed with Adobe Photoshop CS5 software (Adobe Systems, San Jose, CA, USA) using only contrast optimization, Gaussian and channel overlay functions affecting the whole image equally.

The quality of the mitotic chromosome spreads was evaluated based on chromosome morphology, the absence of overlapping chromosomes, and the clarity of the spread from any debris on the slide. Chromosome spreads were considered high quality if they exhibited well-delineated, intact chromosomes with minimal background noise, which allowed for easy and accurate identification of individual chromosomes eventually facilitating accurate FISH signal detection. Metaphase index was calculated as the percentage of cells at metaphase stage. Based on the quality of the chromosome preparations and the highest mitotic indices, the best synchronization method was selected and tested for suitability for wild *Crocus* accessions, using two *C. cartwrightianus* accessions as use cases.

### Probe labeling and fluorescent *in situ* hybridization

The probe “18SrRNAgene_Bv_probe1” [30] was used for the detection of the rDNA and was labeled with biotin-11-dUTP (Dyomics) by PCR and detected by Streptavidin-Cy3 (Sigma-Aldrich). The probe pXV1 [31] for the 5S rRNA gene was labeled with digoxygenin-11-dUTP by PCR and detected by anti-digoxigenin–fluorescein isothiocyanate (FITC; both from Roche Diagnostics). The hybridization procedure was performed as described previously [31]. Chromosomes were counterstained with DAPI (Honeywell, Charlotte, NC, USA). Prior to FISH, according to the amount of cytoplasm visible under light microscope, we pre-treated the slides with 100 μg/ ml RNase in 2× SSC for 30 min, followed by 200 μl of 10 μg/ ml pepsin in 10 mM HCl for 15 to 30 min.

## Results and Discussion

### Ice water pretreatment is most effective for obtaining metaphase spreads of high-quality

The four methods introduced above (Hydroxyurea-colchicine (HC), Nitrous Oxide (NO), Hydroxyquinoline (HQ), and Ice Water (IW)) were compared for their effectiveness in obtaining properly dispersed mitotic chromosomes of *C. sativus*, focusing on improving the quality of spreads and their suitability for subsequent FISH analysis. These methods were applied to root tips of *C. sativus* and *C. cartwrightianus* species. In total, 22,507 nuclei were evaluated and representative metaphases were selected for illustration (Figure 1).

**Figure 1:**
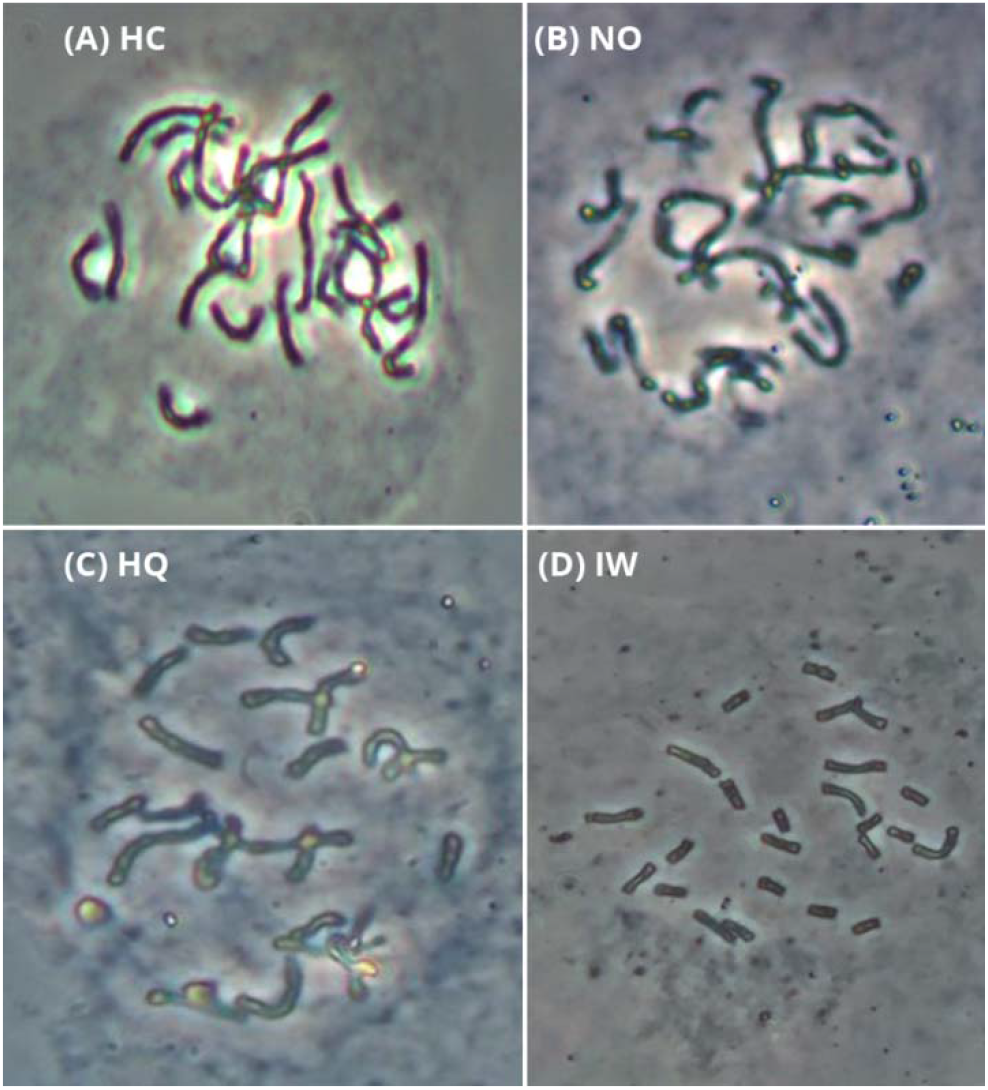
Exemplary mitotic chromosome spreads of *C. sativus*, obtained with each of the four treatments: HC (A), NO (B), HQ (C), and IW (D). The spreads shown in this figure were taken by phase contrast microscope (Zeiss Axioskop 40) at a magnification power of 400x.

Comparing all four methods, HC (Figure 1A, Table 2) yielded the lowest metaphase index (1.08%). HC-derived metaphases usually featured chromosomes that were difficult to count, were unrecognizable, and had much overlapping of chromosome arms. Hence, using HC for chromosome preparation and FISH analysis of *Crocus* is not recommended. The same trend was observed for the NO and HQ (Figure 1B-1C, Table 2).

**Table 2:**
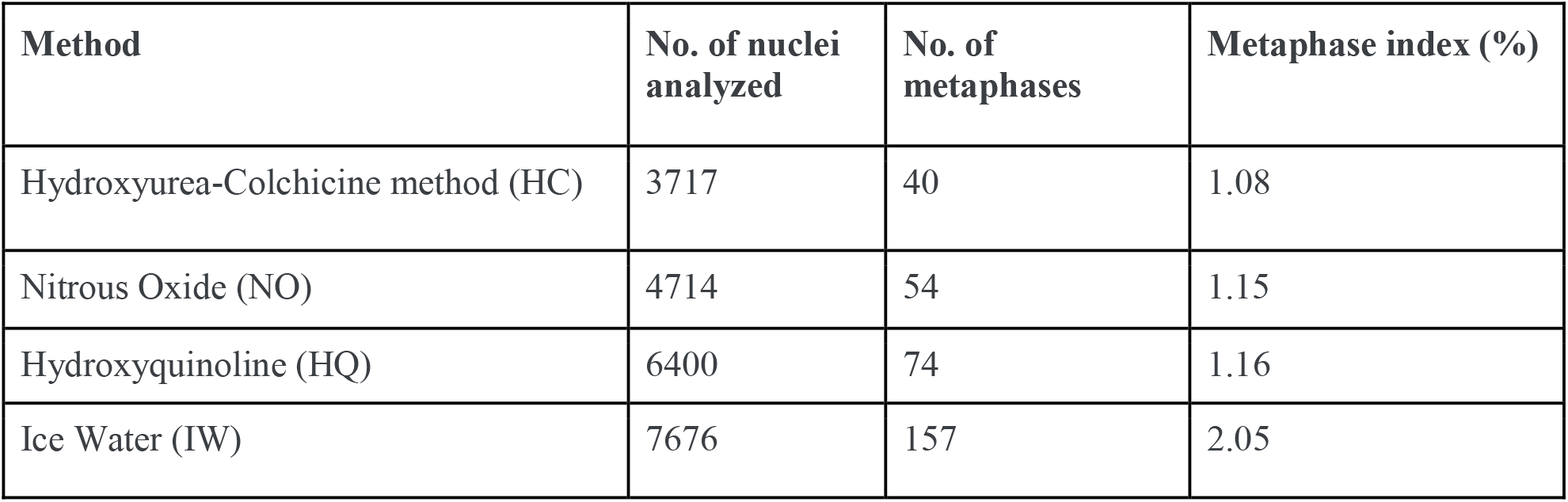
Comparison of mitotic indices in four methods used for chromosome preparation from *C. sativus* materials.

Similar to HC, the Nitrous Oxide (NO) and Hydroxyquinoline (HQ) methods showed limited effectiveness in producing high-quality chromosome spreads. NO-prepared spreads (Figure 1B) often resulted in chromosomes that were difficult to distinguish, with irregular morphology and overlapping chromosome arms, complicating accurate identification and analysis. The HQ method (Figure 1C, Table 2) yielded comparable results, with a metaphase index of 1.16%, slightly higher than NO but still insufficient for obtaining consistently clear, non-overlapping chromosomes. Weighing all four methods against each other, IW yielded the highest metaphase index (2.05%). We conclude that this pretreatment arrested the cells in metaphase twice more often than the other three methods (Table 2). Moreover, this method is preferred in terms of chromosome morphology, if the chromosomes are to be counted or analyzed by FISH procedure, as they were of preferred shape and length with no overlapping (Figure 1D).

### Ice water pretreatment is also effective for wild crocus species and allows following the nuclei through mitosis

To test the most effective method also for wild *Crocus* species, we applied the ice water pretreatment also to *C. cartwrightianus*, the diploid progenitor species of *C. sativus*. Using IW pretreatment, nuclei from both species, *C. sativus* and *C. cartwrightianus*, were easily stained with DAPI, resulting in clear, well-resolved chromosome spreads that allowed for detailed visualization of each stage of mitosis (Figure 2). The preparations showed high-quality chromosome morphology, with distinct and non-overlapping chromosomes across the mitotic phases, facilitating accurate structural analysis as shown later by the FISH analysis. In the DAPI-stained images of both *C. sativus* and *C. cartwrightianus* prepared using the IW method, all stages of mitosis (interphase, prophase, prometaphase, metaphase, anaphase, and telophase) were clearly observed. Each stage displayed its characteristic features of chromosome condensation and organization (Figure 2).

**Figure 2:**
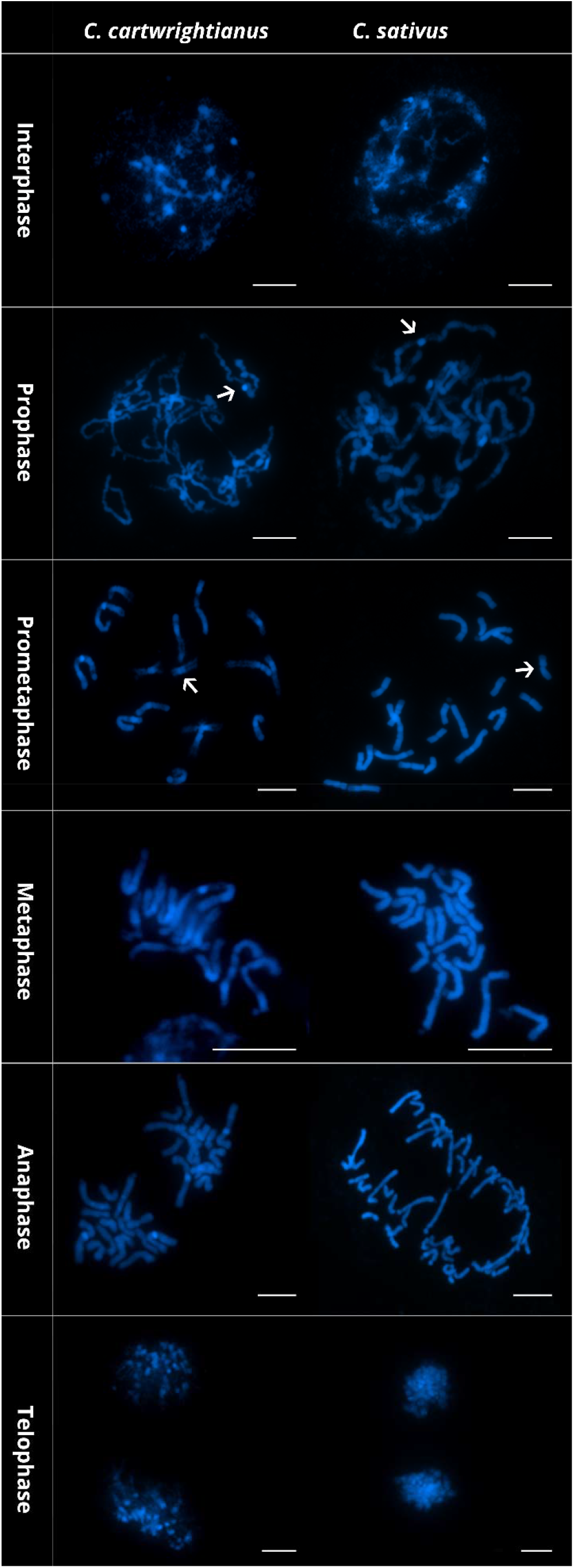
DAPI-stained chromosome spreads of *C. cartwrightianus* and *C. sativus*. The scale bar equals 10 μm. The arrow indicates the centromeric constriction.

The DAPI-staining of both *C. sativus* and *C. cartwrightianus* chromosomes enabled the identification of intercalary and terminal heterochromatin and of the mostly weakly stained centromeres, often also visible as a constriction (Figure 2; arrowed).

### Metaphases resulting from ice water pretreatment are well-suited for FISH follow-up

Chromosome spreads obtained from root tips prepared by the IW method were evaluated for their applicability for FISH analysis using 18S-5.8S-25S and 5S rDNA probes (Figure 3).

**Figure 3:**
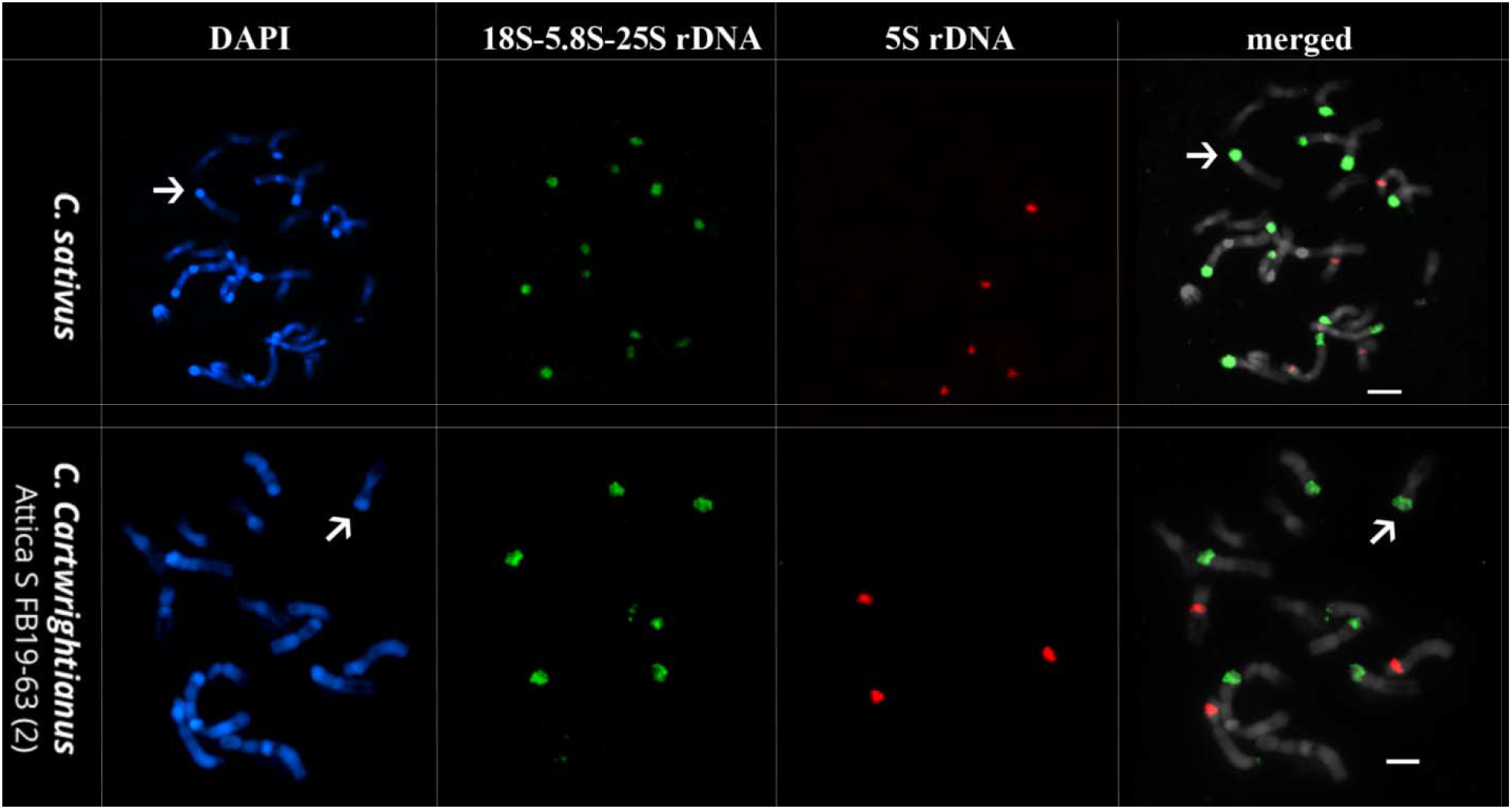
Fluorescent *in situ* hybridization (FISH) of *C. sativus* and *C. cartwrightianus* (name given in each panel). DAPI-stained mitotic chromosomes are shown in blue. Probes used are 18S-5.8S-25S rRNA genes (green), and 5S rRNA genes (red).

As presented (Figure 3), the chromosome spreads produced with IW were highly useful for follow-up analysis by FISH. In this example, the genetic identity can be assigned to the large terminal chromosome regions, which are strongly condensed and detectable as DAPI-positive blocks. These consist mostly of rRNA genes (Figure 3, arrowed, Table 3). In *C. sativus*, we detected 12 arrays of 18S-5.8S-25S rRNA genes, including five major sites and seven moderate sites (Figure 3, arrowed), and five major 5S rRNA gene arrays (Figure 3, arrowed). Eight hybridization signals were detected in metaphase chromosomes of *C. cartwrightianus* using 18S-8S-25S rDNA probe, of which six were strong signals while the remaining two were weak signals. FISH analysis using 5S rDNA revealed five moderate signals in *C. sativus* and three strong signals in *C. cartwrightianus*. The number, relative signal strength and presence on chromosomes of the two rDNA sites are listed in Table 3.

**Table 3:**
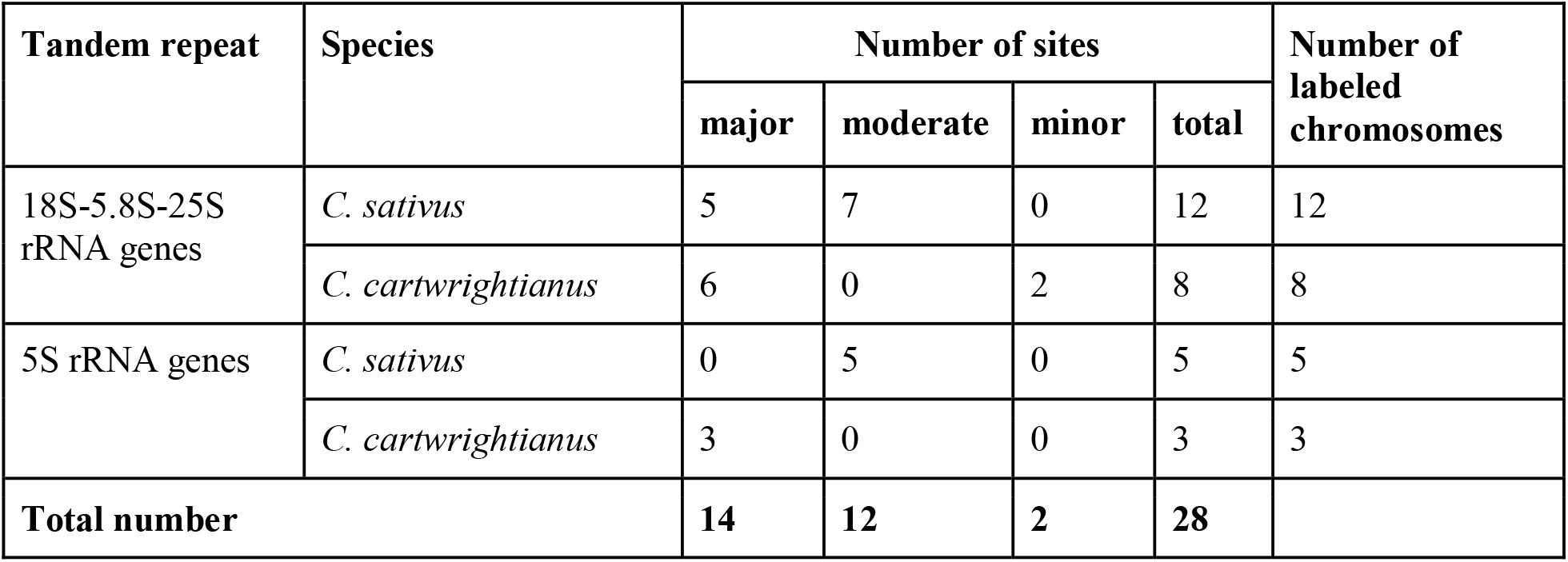
Hybridization of cytogenetic landmarks in *C. sativus* and *C. cartwrightianus* using two rDNA probes.

| Tandem repeat | Species | Number of sites |  |  |  | Number of labeled chromosomes |
| --- | --- | --- | --- | --- | --- | --- |
|  |  | major | moderate | minor | total |  |
| 18S-5.8S-25S rRNA genes | *C. sativus* | 5 | 7 | 0 | 12 | 12 |
|  | *C. cartwrightianus* | 6 | 0 | 2 | 8 | 8 |
| 5S rRNA genes | *C. sativus* | 0 | 5 | 0 | 5 | 5 |
|  | *C. cartwrightianus* | 3 | 0 | 0 | 3 | 3 |
| **Total number** |  | **14** | **12** | **2** | **28** |  |

## Conclusion

Summarizing, we present a comprehensive comparison of four methods (HC, NO, HQ, and IW) for preparing mitotic chromosome spreads in *Crocus sativus*. The results indicate that ice water pretreatment (IW) is more suitable, yielding the highest metaphase index (2.05%) and providing the best chromosome morphology for further analysis. In contrast, HC, NO, and HQ yielded lower metaphase indices and produced chromosomes with poor morphology, making them less suitable for cytogenetic studies. IW pretreatment was also equally well with *C. cartwrightianus*, the diploid wild progenitor species.

The suitability of this method for cytogenetic analysis across different *Crocus* species was demonstrated by a) enabling us of following the different stages of mitosis in both species with a clear identification of DAPI-stained metaphase chromosomes that provided detailed insights into chromosomal structure, b) the successful FISH hybridization to chromosomes enabling the visualization of key chromosomal features such as heterochromatin and centromeres in both cultivated and wild crocus species.

## Abbreviations

DAPI: 4,6-diamidino-2-phenylindole
FISH: Fluorescence *in situ* hybridization
Hq: 8-hydroxyquinoline
rDNA: Ribosomal DNA
RT: Room temperature
DW: Distilled water
SSC: Saline sodium citrate

## Acknowledgments

We are grateful to Frank Blattner and Dörte Harpke (IPK Gatersleben, Gatersleben, Germany) for providing some of the used plant materials.

## Author Contributions

AE and TH designed the study. AE and SL selected and cultivated the plants. AE, SL and SB performed the slide preparations and the FISH hybridisation experiments. AE analyzed the data and prepared the figures and tables. AE and TH wrote the manuscript. All authors read and approved the final manuscript.

## Availability of data and materials

Not applicable.

## Ethics approval and consent to participate

Not applicable.

## Consent for publication

Not applicable.

